# The 5-LOX/COX-2 cross-over metabolite, hemiketal E_2_, enhances VEGFR2 activation and promotes angiogenesis

**DOI:** 10.1101/2022.05.13.491890

**Authors:** Fumie Nakashima, Juan A. Giménez-Bastida, Paula B. Luis, Sai H. Presley, Robert E. Boer, Manuel Chiusa, Takahiro Shibata, Gary A. Sulikowski, Ambra Pozzi, Claus Schneider

## Abstract

Consecutive oxygenation of arachidonic acid by 5-lipoxygenase (5-LOX) and cyclooxygenase-2 (COX-2) yields the hemiketal (HK) eicosanoids, HKE_2_ and HKD_2_. HKs stimulate angiogenesis by inducing endothelial cell tubulogenesis in culture; however, how this process is regulated has not been determined. Here, we identify vascular endothelial growth factor receptor 2 (VEGFR2) as a mediator of HKE_2_-induced angiogenesis *in vitro* and *in vivo*. HKE_2_ treatment of human umbilical vein endothelial cells dose-dependently increased phosphorylation of VEGFR2 and the downstream kinases ERK and Akt that mediated endothelial cell tubulogenesis. *In vivo*, HKE_2_ induced the growth of blood vessels into polyacetal sponges implanted in mice. HKE_2_-mediated effects *in vitro* and *in vivo* were blocked by the VEGFR2 inhibitor vatalanib, indicating that the pro-angiogenic effect of HKE_2_ was mediated by VEGFR2. We found that HKE_2_ covalently bound and inhibited PTP1B, a protein tyrosine phosphatase that dephosphorylates VEGFR2, thereby providing a possible molecular mechanism for how HKE_2_ induced pro-angiogenic signaling. Our studies indicate that biosynthetic cross-over of the 5-LOX and COX-2 pathways gives rise to a potent lipid autacoid that regulates endothelial cell function *in vitro* and *in vivo*.

**Significance:** Angiogenesis, the growth of new blood vessels from existing vessels, contributes to both physiological and pathological conditions, including tissue repair after injury and tumorigenesis. Novel approaches to control pathologic angiogenesis are urgently needed since current therapy targeting the pro-angiogenic receptor VEGFR2 has significant side effects. We show that a metabolite of arachidonic acid, formed in a biosynthetic cross-over of the enzymes that generate the pro-inflammatory leukotriene and prostaglandin mediators, respectively, promotes VEGFR2 activation to induce angiogenesis. This finding suggests that common drugs targeting the arachidonic acid pathway may be viewed as valid anti-angiogenic candidates.

## Introduction

Arachidonic acid is the common substrate for the leukotriene and prostaglandin biosynthetic pathways. Following the initial oxidation of arachidonic acid by 5-lipoxygenase (5-LOX) or cyclooxygenase-2 (COX-2), respectively, the biosynthetic reactions proceed along separate pathways (1-3). Arachidonic acid can also undergo consecutive oxygenation by 5-LOX and COX-2 to yield two hemiketal (HK) eicosanoids, HKE_2_ and HKD_2_ (4) as well as the 5-hydroxyprostaglandins, 5-OH-PGE_2_ and 5-OH-PGD_2_ (5). The cross-over biosynthetic pathway entails initial oxygenation of arachidonic acid by 5-LOX to form 5*S*-HETE that subsequently undergoes oxygenation by COX-2. In HK biosynthesis COX-2 reacts 5*S*-HETE with three molecules of oxygen to form a di-endoperoxide as the unstable enzymatic product (6). Spontaneous rearrangement of the peroxide bonds of the di-endoperoxide yields HKE_2_ and HKD_2_ as stable products (Fig. 1) (4). Oxygenation of 5-HETE is selective for the natural, 5-LOX derived 5*S*-enantiomer, and a specific reaction of COX-2 whereas COX-1 does not react with either 5*S*-nor 5*R*-HETE (6).

**Fig. 1.**
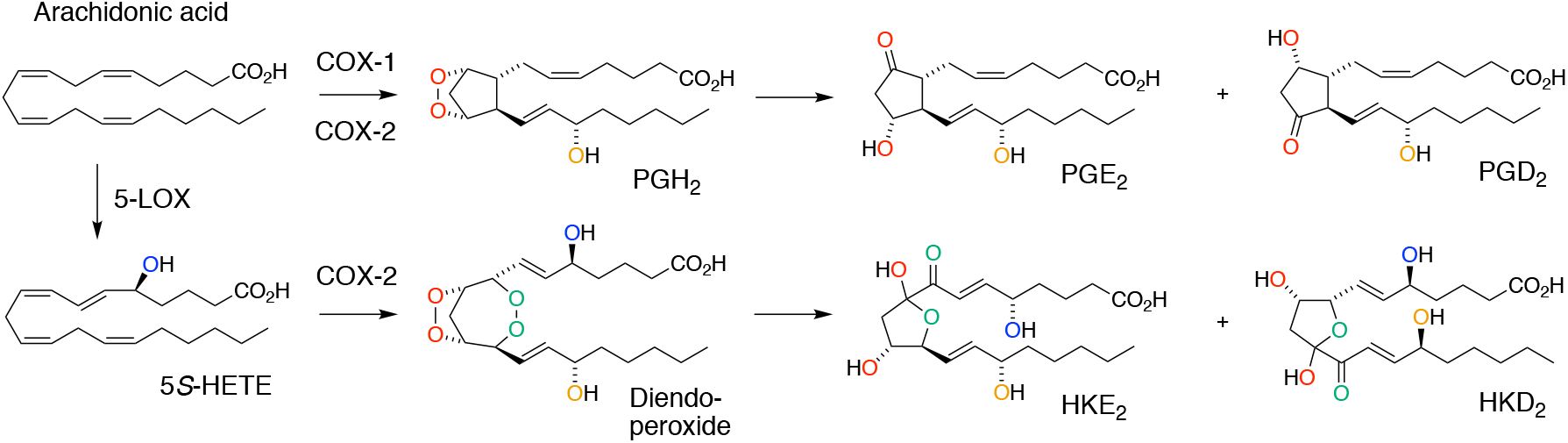
The 5-LOX/COX-2 cross-over biosynthetic pathway to hemiketal (HK) eicosanoids HKE_2_ and HKD_2_ in comparison to prostaglandin formation. The top reactions show prostaglandin biosynthesis by COX-1 and COX-2. The prostaglandin endoperoxide PGH_2_ formed in the COX reaction undergoes rearrangement to PGE_2_ and PGD_2_. The reactions below show the cross-over pathway initiated by 5-LOX formation of 5*S*-hydroxyeicosatetraenoic acid (5*S*-HETE) from arachidonic acid followed by COX-2 catalyzed formation of a di-endoperoxide that rearranges to HKE_2_ and HKD_2_.

The expression of 5-LOX and COX-2 by activated neutrophils and macrophages, respectively, suggested mixtures of these cells as a source of HK biosynthesis (7). Freshly isolated human leukocytes, when activated with bacterial lipopolysaccharide (LPS) and calcium ionophore A23187, robustly formed HKE_2_ and HKD_2_, and this was blocked by inhibitors of 5-LOX or COX-2, respectively (8). Levels of HKs were around 1% relative to LTB_4_ and about equal to PGE_2_ when leukocyte mixtures were activated with LPS for less than 2 h. The time course of HK formation paralleled the transient availability of the 5-HETE substrate rather than the steadily increasing expression of COX-2 over longer activation times with LPS (8). This suggested that HKs might be formed early during an inflammatory event *in vivo*, when the response is dominated by 5-LOX-expressing neutrophils prior to influx of activated macrophages that express COX-2 (9).

Little is known about the biological activities of HKE_2_ and HKD_2_ besides their ability to stimulate *in vitro* tubulogenesis of vascular endothelial cells and to inhibit platelet aggregation (4, 10). Lack of a ready supply of HKs has hindered their biological characterization. HKs are cumbersome to produce using enzymatic synthesis (11), and their total chemical synthesis was not completed until recently (10). With HKE_2_ available by chemical synthesis, we have analyzed signaling pathways activated by HKE_2_ with emphasis on vascular endothelial cell tubulogenesis and proangiogenic effects *in vivo*.

## Results

### HKE_2_ promotes tyrosine phosphorylation in human endothelial cells

We analyzed global protein tyrosine phosphorylation in human vascular endothelial cells (HUVEC) treated with HKE_2_. A time course experiment of cells treated with HKE_2_ (500 nM) for 30-120 min showed highest tyrosine phosphorylation at 30 min (Fig. 2A). When HUVEC were treated with different concentrations of HKE_2_ (0-500 nM) for 30 min, 100 nM HKE_2_ had the strongest effect while at 500 nM HKE_2_ the effect was still evident although less than with 100 nM HKE_2_ (Fig. 2B). The potent induction of protein tyrosine phosphorylation suggested involvement of a cell surface receptor in response to treatment with HKE_2_. Since receptor tyrosine kinases (RTK) are known to be involved in angiogenesis (12, 13), we screened RTK activation by HKE_2_ using an RTK phosphorylation array. HUVEC were treated with HKE_2_ at 50, 100, and 500 nM for 30 min (Fig. 2C). Out of 49 RTK analyzed, ALK, EphA10, RYK, Tie1, and VEGFR2 were consistently phosphorylated in two independent experiments (Fig. 2 D). VEGFR2 was selected for further analysis due to its established role in mediating endothelial cell angiogenesis (14, 15) and as a receptor regulated by arachidonic acid-derived lipids (16, 17).

**Fig. 2.**
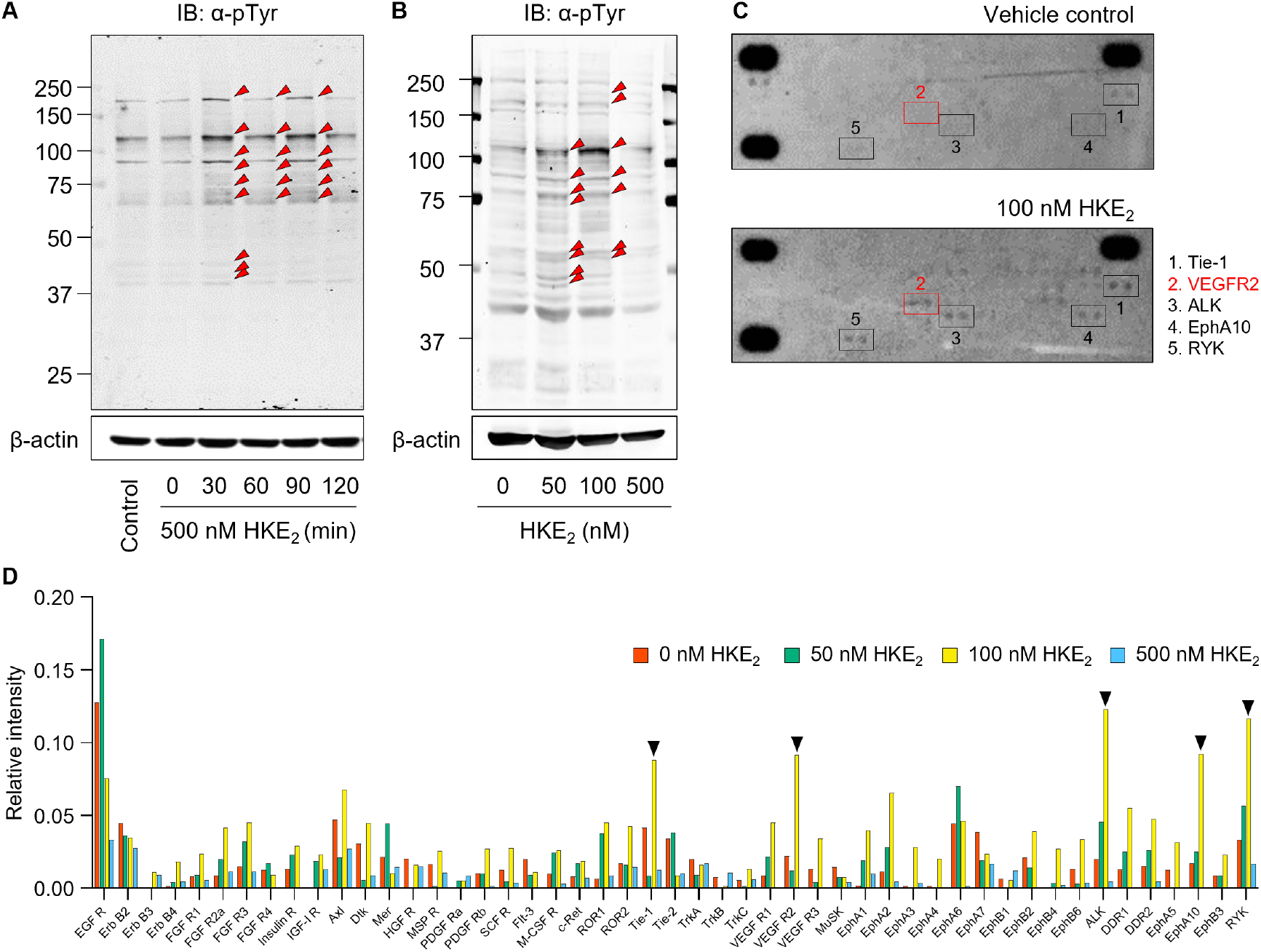
Protein phosphorylation in endothelial cells treated with HKE_2_. HUVEC were treated with (A) 500 nM of HKE_2_ for 0-120 min or (B) 0-500 nM of HKE_2_ for 30 min, and cellular protein was analyzed by SDS-PAGE and immunoblotting using antibodies against phospho-tyrosine (α-pTyr) and β-actin. Red triangles indicate protein bands that showed increased phosphorylation in response to treatment with HKE_2_ compared to vehicle control. (C) A human phospho-Receptor Tyrosine Kinase (RTK) array was used to analyze HKE_2_-induced phosphorylation of 49 RTKs in HUVEC lysates. X-ray film images of the membranes treated with vehicle or 100 nM HKE_2_ are shown. Protein spots that showed differences in intensity between HKE_2_ and vehicle control are boxed. (D) Quantification of the RTK array results using ImageJ software. The five RTK boxed in panel (C) are marked with a black triangle. One of two independent repeats of the RTK array using 0-500 nM HKE_2_ giving similar results is shown.

### HKE_2_ promotes VEGFR2 phosphorylation and downstream signaling

Western blot analysis of HUVEC treated with HKE_2_ showed dose-dependent phosphorylation of VEGFR2 at Y1175, one of the major autophosphorylation sites, consistent with the results from the RTK array (Fig. 3A). The effect of HKE_2_ was similar to the activation achieved by VEGF_165_ that was used a positive control (Fig. 3B). Phosphorylation of VEGFR2 induced by HKE_2_ or VEGF_165_ was decreased by pretreatment of HUVEC with vatalanib, a selective inhibitor of VEGFR2 kinase activity (Fig. 3B) (18, 19). Vatalanib also prevented HKE_2_- and VEGF_165_-induced phosphorylation of downstream ERK and Akt (Fig. 3C and D), two kinases known to be activated upon phosphorylation of VEGFR2 at Y1175 (20, 21).

**Fig. 3.**
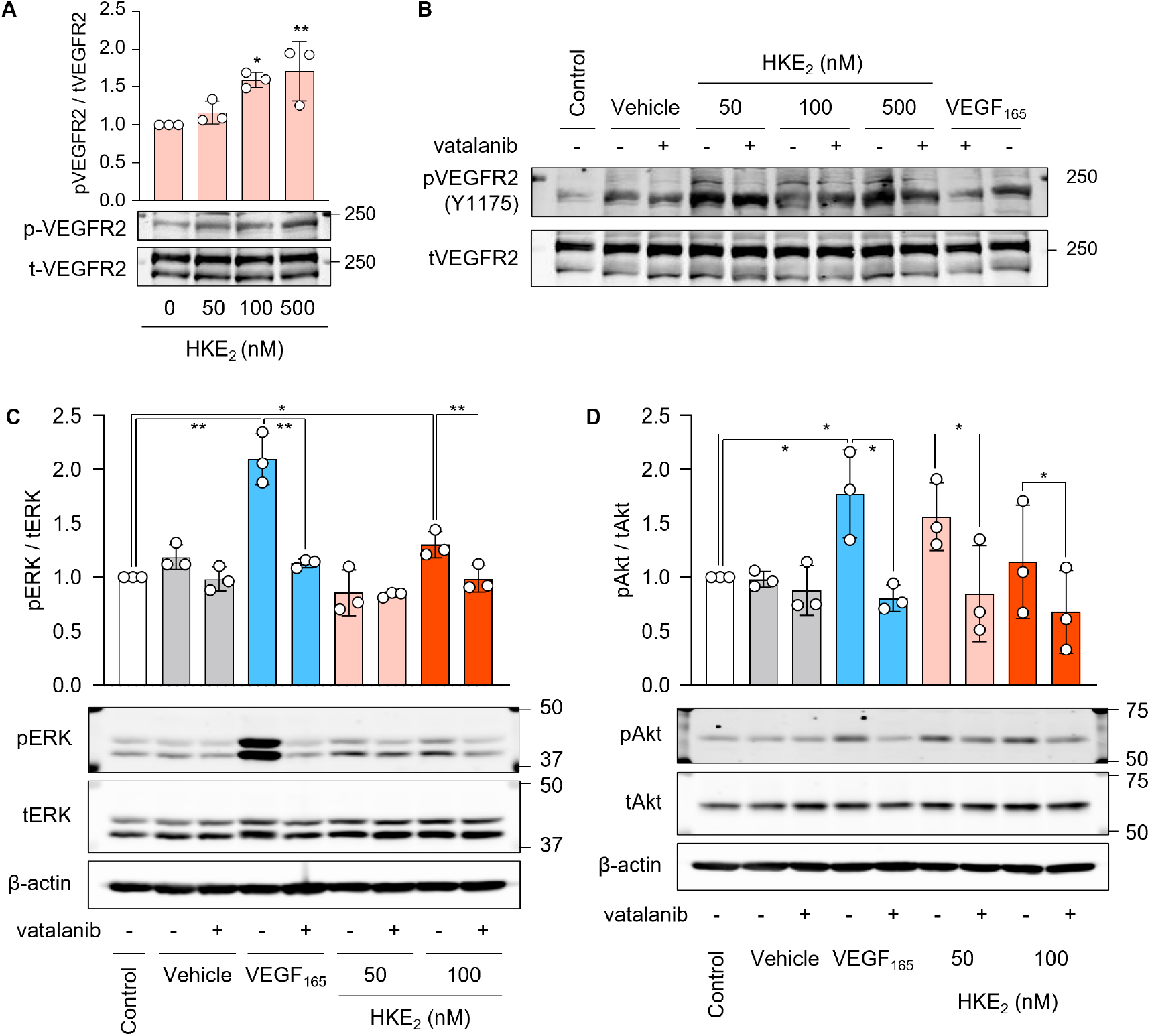
Phosphorylation of VEGFR2 and downstream kinases in endothelial cells treated with HKE_2_. (A) Western blots of phosphorylated and total VEGFR2 in cell lysates of HUVEC treated with 0-500 nM HKE_2_ for 30 min. The graph shows the ratio of phosphorylated to total VEGFR2 from n = 3 independent repeats. Data are presented as means ±S.D. (*) indicates significant differences (*; *p < 0*.*05*, **; *p < 0*.*01*) between vehicle (0 nM HKE_2_) and HKE_2_-treated cells. (B) Cells were treated with the VEGFR2 inhibitor vatalanib 30 min prior to HKE_2_ treatment, and phosphorylation of VEGFR2 was analyzed by Western blotting. (C, D) HUVEC were incubated for 1 h with or without vatalanib, and then treated with vehicle, VEGF_165_ (50 ng/mL) or the indicated concentration of HKE_2_. Cell lysates were prepared and subjected to Western blotting for phosphorylated and total ERK and Akt, respectively, and for β-actin. Bar graphs show the ratio of phosphorylated over total proteins obtained from n = 3 independent repeats. Data are presented as means ±S.D. (*) indicates significant differences (*; *p < 0*.*05*, **; *p < 0*.*01*).

### HKE_2_ stimulates tubulogenesis through VEGFR2 activation

HUVEC were treated with HKE_2_, VEGF_165_, or vehicle for 6 h with or without vatalanib pretreatment, and cellular network formation was determined using a matrigel based angiogenesis assay (Fig. 4A). Cells treated with 50 nM HKE_2_ showed a significant increase in characteristic features of cellular tubulogenesis in vitro that were related to forming connections between cells and ring-like structures (Fig. 4B-G). The effects of HKE_2_ were similar to treatment with VEGF_165_. All of the tubulogenic parameters increased by HKE_2_ were decreased by pretreatment with vatalanib. An increase in tubulogenesis appeared visually evident also at the lowest concentration of HKE_2_ used (10 nM) but the quantitative analysis showed no significant effect compared to vehicle control.

**Fig. 4.**
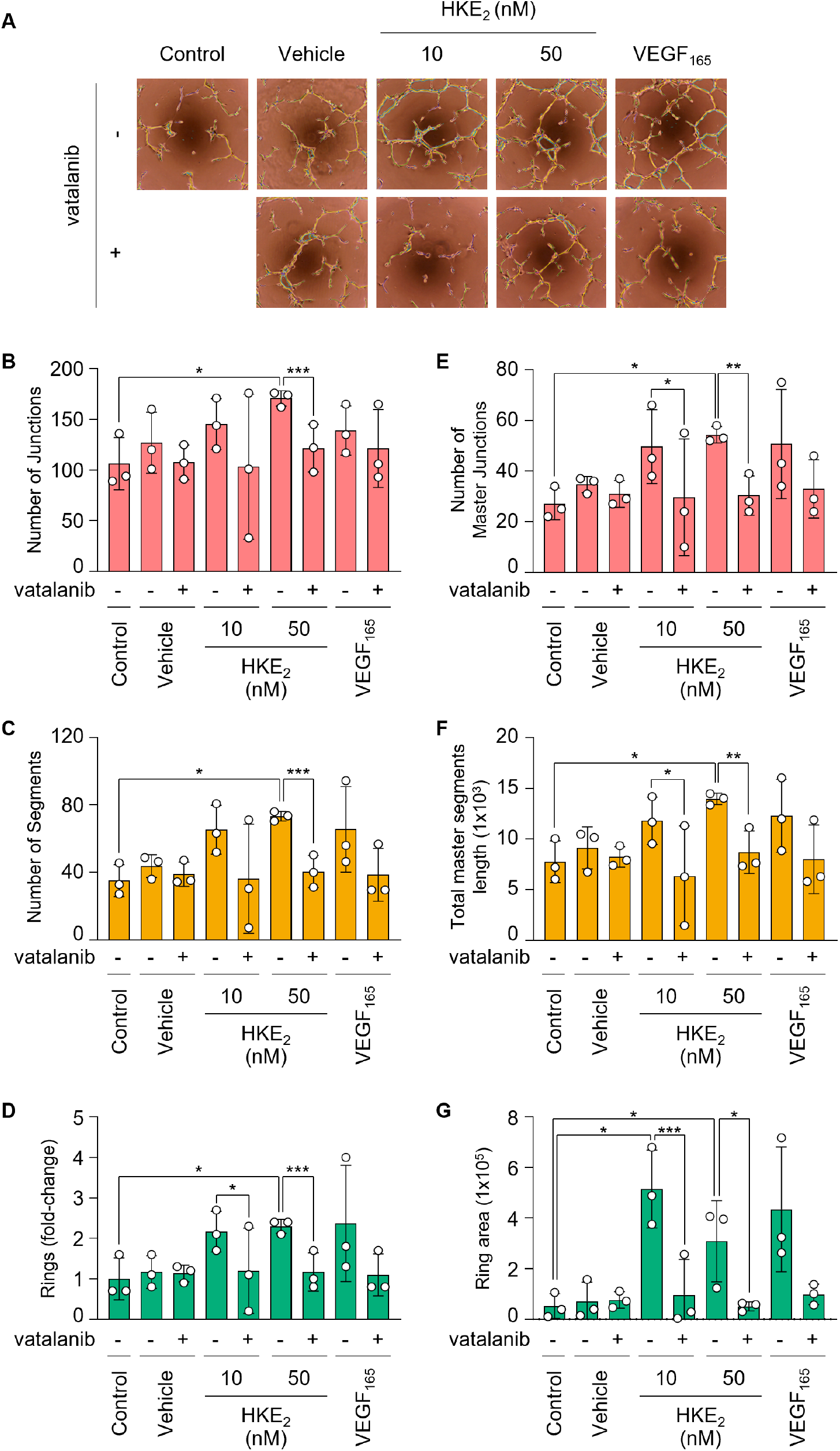
HKE_2_ induces formation of capillary-like structures by endothelial cells. (A) HUVEC were placed in serum-free Medium 200PRF onto Matrigel coated 96-well plates in the absence or presence of the VEGFR2 inhibitor, vatalanib. After 2 h incubation HKE_2_ or VEGF_165_ (50 ng/mL) was added to the wells. Representative images of tube-like structures were taken 6 h after HKE_2_ treatment. (B-G) Capillary network formation was quantified using ImageJ software with the Angiogenesis Analyzer plugin counting the number of (B) junctions, (C) segments (D) rings, (E) master junctions, (F) length of total master segments, and (G) ring area from two independent experiments. Data are presented as means ±S.D. (*) indicates significant differences (*; *p < 0*.*05*, **; *p < 0*.*01*, ***; *p < 0*.*005*).

### Kinase inhibitors decrease HKE_2_-induced HUVEC tubulogenesis

HUVEC were treated with the MAPK/ERK inhibitor U0126 or the PI3 kinase inhibitor LY294002 for 30 min prior to HKE2 stimulation, and cellular network formation was determined after 6 h. The ERK inhibitor significantly decreased the formation of cellular junctions, segments, and rings while inhibition by the PI3 kinase inhibitor was significant only for the decrease in cellular junctions and segments (Fig. 5). Inhibition of HKE2-induced tubulogenesis by the ERK and PI3 kinase/Akt signaling pathways inhibitors confirmed a role of these signaling pathways in mediating tubulogenesis downstream of VEGFR2 activation (Fig. 3C and D).

**Fig. 5.**
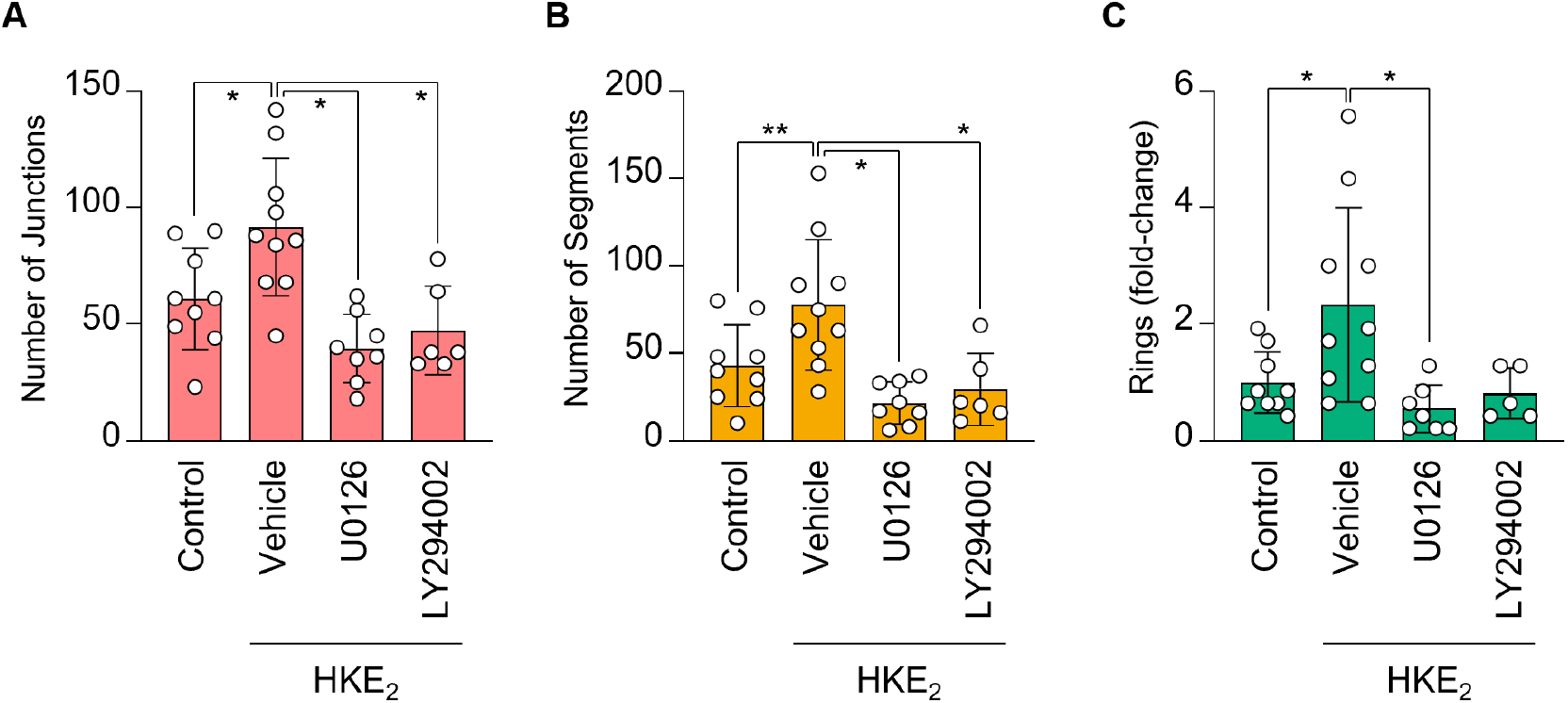
Kinase inhibition in HKE_2_-induced tubulogenesis of HUVEC. HUVEC were plated in 96-well plates containing reduced growth factor Matrigel and were treated with vehicle (DMSO), the MAPK/ERK inhibitor U0126 (10 µM) or the PI3 kinase inhibitor LY294002 (1 µM). After 30 minutes, cells were treated with ethanol (vehicle) or HKE_2_ (500 nM). After 6 hours, capillary network formation was visualized and quantified by ImageJ software with the Angiogenesis Analyzer plugin counting the number of (A) junctions, (B) segments, and (C) rings. Data are representative of 3 independent experiments. Data are presented as means ±S.D. (*) indicates significant differences (*; *p < 0*.*05*, **; *p < 0*.*01*).

### PGE_2_ and 5*S*-HETE do not stimulate HUVEC tubulogenesis

The effect of HKE_2_ was compared to 5*S*-HETE as the 5-LOX derived substrate in HK biosynthesis by COX-2 (6) and to PGE_2_ as a COX-2 product with reported pro-angiogenic properties (22-24). Compared to vehicle treated cells, PGE_2_ did not change the number of junctions, segments, or rings formed by HUVEC when used at 100 nM, a concentration ∼10-fold above the EC_50_ required for activation of its receptors (Supplementary Fig. S1). Similarly, 5*S*-HETE at 100 nM did not stimulate tubulogenesis (Supplementary Fig. S1). Thus, consecutive oxygenation of arachidonic acid by 5-LOX and COX-2 yields HKE_2_ as an eicosanoid with potent pro-tubulogenic activity that was not matched by the biosynthetic intermediate 5*S*-HETE or by the prototypical COX-2 product, PGE_2_.

### HKE_2_ induces angiogenesis *in vivo*

To test whether HKE_2_ has pro-angiogenic activity *in vivo* a subcutaneous sponge assay was performed as described previously (25). Sterile polyvinyl acetal round sponges were implanted subcutaneously in the right and left side of the back of adult female mice. The sponges were injected every other day with 50 μL of vehicle (coconut medium-chain triglyceride (MCT) oil), HKE_2_ (10 µM in MCT oil), or VEGF165 (5 μg/mL in PBS). In parallel, animals received vatalanib (50 mg/kg) or vehicle (DMSO/polyethylene glycol 400/0.9% sodium chloride: 5/47.5/47.5) by oral gavage daily starting from the day of the first injection of HKE_2_. After two weeks, mice were injected prior to sacrifice with rhodamine-dextran for visualization of blood vessels within sponges. Sponges were subsequently collected and analyzed under an epifluorescence microscope. Visual inspection of the sponges showed high vascularization with HKE_2_ treatment that was similar to that observed in sponges injected with VEGF165 and near absent in animals receiving MCT oil only (Fig. 6A and Supplementary Fig. S2). Co-treatment with vatalanib decreased vascularization in animals receiving HKE_2_ or VEGF_165_, while no effect was observed in inhibitor-treated mice receiving MCT oil. Quantitative analysis of stained vessels by microscopy confirmed the observation made by visual inspection. Vascularity within sponges was expressed as a percentage of area occupied by rhodamine-dextran-positive structures per microscopic field. Quantification showed a significant increase of blood vessel formation by both HKE_2_ and VEGF_165_, and angiogenesis induced by both agents was significantly decreased by treatment with vatalanib (Fig. 6B).

**Fig. 6.**
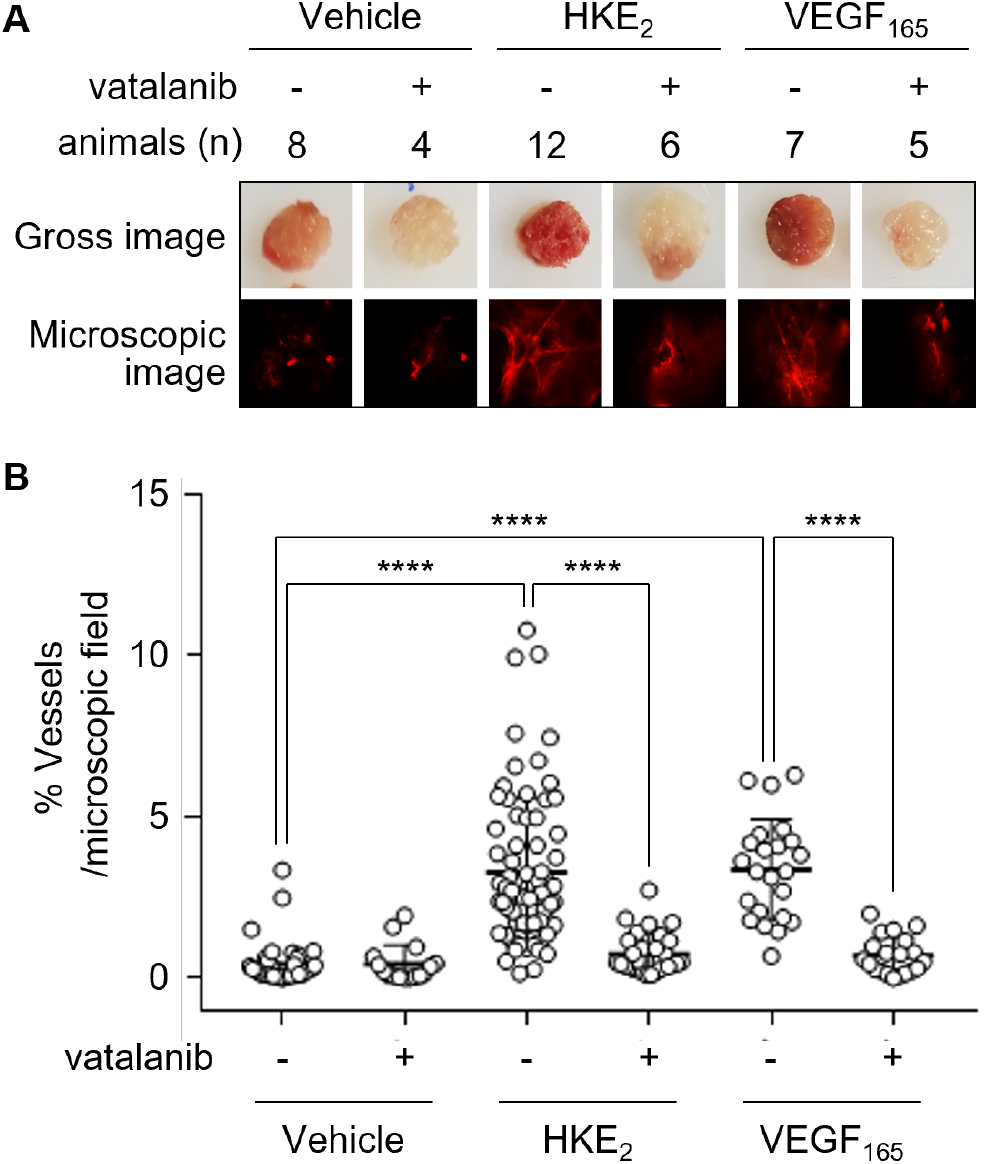
*In vivo* angiogenesis induced by HKE_2_. (A) Gross and microscopic images of polyvinyl acetal CF-50 sponges implanted under the dorsal skin of BALB\c female mice (12 weeks of age, 20 g). The sponges were injected every other day for 2 weeks with 50 μL of vehicle (coconut medium-chain triglyceride (MCT) oil, n = 8 animals), HKE_2_ (10 µM in MCT oil, n = 12), or VEGF_165_ (5 μg/mL in PBS, n = 7). In addition, animals received vatalanib (50 mg/kg) or vehicle (DMSO/polyethylene glycol 400/0.9% sodium chloride: 5/47.5/47.5) by gavage daily (vehicle + inhibitor, n = 4; HKE_2_ + inhibitor, n = 6; VEGF_165_ + inhibitor, n = 5) starting from same day of the first injection of HKE_2_. Fifteen minutes prior sacrifice, mice were injected intravenously with 50 μL of rhodamine-dextran (2% in PBS). Sponges were then removed, sectioned, and placed under a fluorescence microscope to visualize vascularization. (B) Vascularity within sponges was quantified by calculating the area occupied by rhodamine-positive structures per microscopic field. The horizontal bar indicates the mean ± S.D. calculated for five-six images/sponge per treatment. Differences were analyzed by ANOVA followed by Dunnett’s multiple comparison analysis. (*) indicates significant differences (****; *p < 0*.*0001*).

### HKE_2_ does not bind VEGFR2 or activate MMP9

As potential mechanism for HKE_2_-mediated phosphorylation of VEGFR2 we tested whether HKE_2_ directly binds to the receptor in order to induce autophosphorylation. Binding of HKE_2_ to the soluble extracellular ligand binding domain of human VEGFR2 was not observed when tested by using microscale thermophoresis (Supplementary Fig. S3), thus arguing against direct activation of VEGFR2 by HKE_2_. We next tested whether HKE_2_ was able to activate matrix metalloproteinase MMP9 resulting in degradation of extracellular matrix and release of matrix bound VEGF (26, 27). Treatment of HUVEC with HKE_2_ in the presence of the MMP inhibitor GM6001 (28) did not decrease phosphorylation of VEGFR2 (Supplementary Fig. S4), suggesting that HKE_2_ did not activate MMP9 to stimulate the release of the VEGFR2 ligand.

### HKE_2_ covalently binds and inhibits tyrosine phosphatase PTP1B

Since HKE_2_ did not activate VEGFR2 directly nor promotes release of VEGF, we tested whether HKE_2_ reduced dephosphorylation of activated VEGFR2. Protein tyrosine phosphatase 1B (PTP1B) dephosphorylates activated VEGFR2 thereby reducing pro-angiogenic signaling in endothelial cells (29). Thus, inhibition of PTP1B might underlie the persistent activation of VEGFR2 observed in HUVEC treated with HKE_2_. HKE_2_ contains an electrophilic enone moiety that is reactive with cysteine nucleophiles as demonstrated by reaction with glutathione (4), suggesting that covalent adduction of HKE_2_ to PTP1B may inhibit this phosphatase via an established mechanism for small molecule electrophilic inhibitors of PTP1B (30).

In order to test PTP1B as a target of HKE_2_, recombinant human PTP1B was incubated with HKE_2_ or the cysteine-reactive probe *N*-ethyl-maleimide (NEM) as positive control to analyze covalent adduction of the electrophilic compounds to the phosphatase. Unadducted cysteine residues of PTP1B remaining after treatment with HKE_2_ or NEM were probed using biotin-maleimide. A 10- and 90-fold molar excess of HKE_2_ dose-dependently decreased detection of PTP1B by biotin-maleimide indicating that reactive cysteine residues in the enzyme were adducted by HKE_2_ and thus no longer available for reaction with the probe (Fig. 7A, B). In addition, a gel shift assay using cysteine-reactive PEG-PC-maleimide (31), a probe that increases the molecular weight of the adducted protein by 5 kD per adducted cysteine residue, identified six protein bands that were shifted to a higher molecular weight relative to unmodified PTP1B (Fig. 7C). This result was consistent with the reported availability of six reactive cysteine residues in PTP1B (30). The intensity of the shifted bands decreased dose-dependently with increasing concentration of HKE_2_ while the band of unmodified PTP1B increased, indicating covalent adduction of HKE_2_ to cysteine residues of PTP1B (Fig. 7D). A 10- and 90-fold molar excess of HKE_2_ decreased the phosphatase activity by about 25% and 40%, respectively, while the same concentration of NEM resulted in about 80% or near complete inhibition, respectively, of phosphatase activity of PTP1B (Fig. 7E). Although protein adduction by the positive control NEM appeared more prominent than with HKE_2_ (Fig. 7A, C), this did not result in a substantially greater inhibition of the enzymatic activity of PTP1B, suggesting that adduction of PTP1B by HKE_2_ has a greater effect on activity than adduction by the much smaller NEM.

**Fig. 7.**
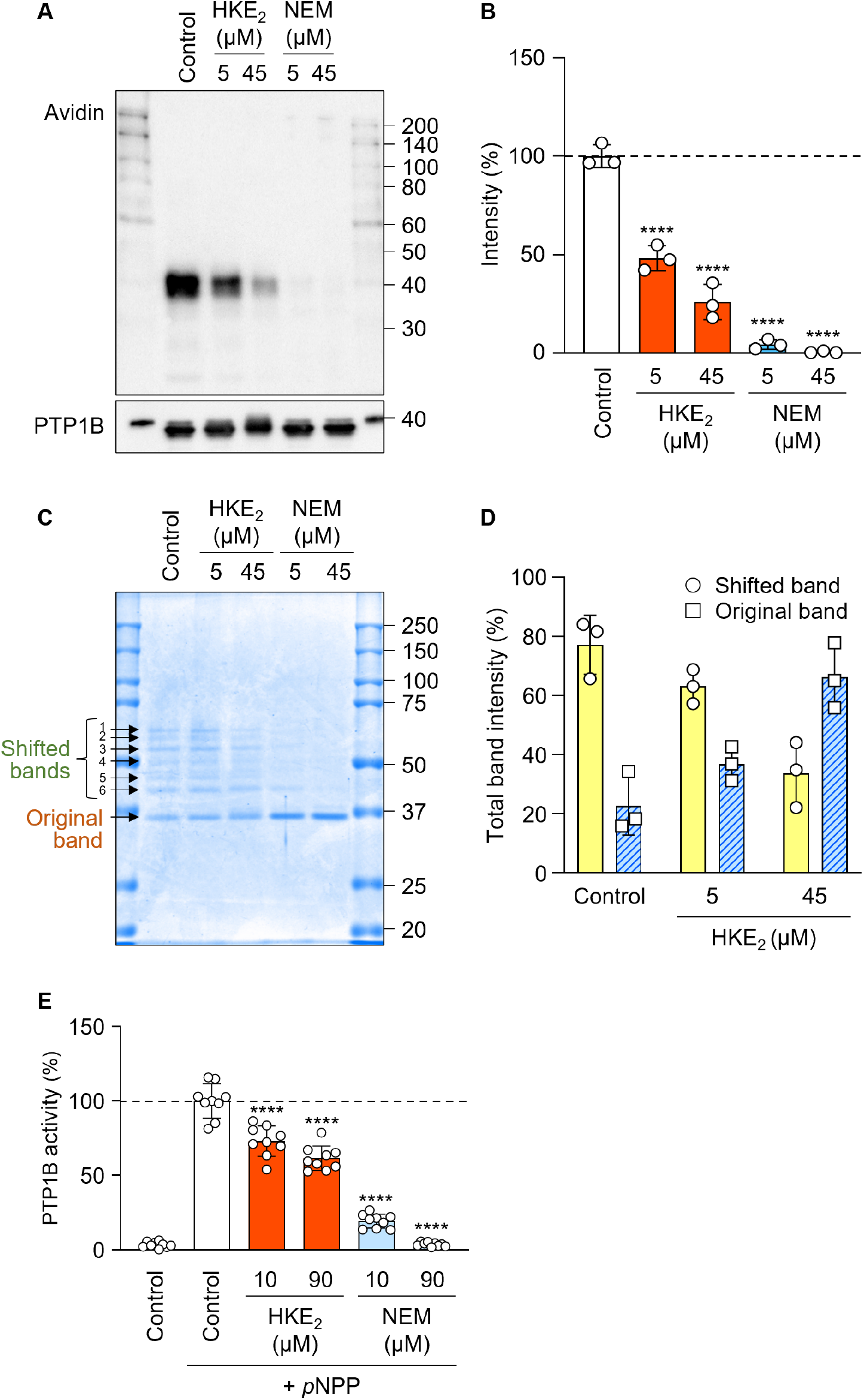
Adduction and inhibition of PTP1B by HKE_2_. (A) Recombinant PTP1B (0.5µM) was treated with HKE_2_ or *N*-ethyl maleimide (NEM) at 5 or 45 µM, respectively, followed by detection of free remaining cysteine residues with biotin-maleimide and analysis by SDS-PAGE. The image shows Western blot detection using HRP-conjugated avidin (top) or an antibody against PTP1B (bottom). (B) The graph shows the ratio of un-modified PTP1B (top panel in A) to total PTP1B (bottom panel in A) from n = 3 independent repeats. Data are presented as means ±S.D. (*) indicates significant differences (****; *p* < 0.0001) between control and HKE_2_ or NEM-treated samples. (C) Recombinant PTP1B (0.5 µM) was treated with HKE_2_ or *N*-ethyl maleimide (NEM) at 5 or 45 µM, respectively, followed by detection of free remaining cysteine residues with PEG-PC-Maleimide and analysis by SDS-PAGE with Coomassie staining. (D) The graph shows the relative abundance of shifted bands (band 1-6) and the original PTP1B band for each treatment from n = 3 independent repeats. (E) Phosphatase activity of PTP1B (1 µM) using *p*NPP as a substrate from n = 3 independent repeats. Data are presented as means ±S.D. (*) indicates significant differences (****; *p* < 0.0001) between control and HKE_2_ or NEM-treated samples.

## Discussion

Eicosanoids formed in each of the three branches of oxygenation of arachidonic acid contribute to the regulation of angiogenesis (32-34). The cyclooxygenase reaction gives rise to prostaglandins and thromboxane, lipoxygenases form hydroxy-eicosatetraenoic acids (HETEs) and leukotrienes, and the cytochrome P450 oxygenation yields epoxide (EET) and terminal hydroxy derivatives of arachidonic acid. EETs from the cytochrome P450 pathway are pro-angiogenic via the induction of growth factors and by increasing the expression of VEGFR2 in endothelial cells. For 8,9-EET the pro-angiogenic activity appears to require further transformation by COX-2 in order to yield the active eicosanoid (17, 35). The lipoxygenase products 15-HETE and 12-HETE have pro-angiogenic properties (36, 37), and the 5-LOX product leukotriene C_4_ was shown to stimulate angiogenesis of endothelial cells in vitro (38). 5-LOX and its 5-HETE metabolite regulated malignant human mesothelial cell survival by inducing expression of VEGF (39) and DNA synthesis and expression of basic fibroblast growth factor 2 in endothelial cells (40), an indication that 5-HETE may be pro-angiogenic. COX-2 has a well-recognized pro-tumorigenic role, and some of the effects are mediated by PGE_2_ via inducing VEGF expression (23, 24) to foster angiogenesis and support tumor growth (41). The role of PGE_2_ as a pro-angiogenic factor is further supported by studies with animals lacking its receptors, e.g., EP2 (42) and EP3 (43, 44)(1).

HKE_2_ is a recently discovered eicosanoid formed in a biosynthetic cross-over of the 5-LOX and COX-2 pathways (Fig. 1) (4). Mixtures of activated human leukocytes formed both HKE_2_ and its isomer HKD_2_ (8), and both HKs activated endothelial cells to form tubular structures in vitro (4). The availability of HKE_2_ (but not HKD_2_) by chemical synthesis (10) has enabled to analyze the mechanism by which HKE_2_ stimulates tubulogenesis *in vitro* and angiogenesis *in vivo*. We identified VEGFR2 as a key mediator when endothelial cells were treated with HKE_2_. Use of the VEGFR2 inhibitor vatalanib abrogated HKE_2_-induced signaling downstream of the receptor and decreased tubulogenesis. In an angiogenesis model *in vivo*, the pro-angiogenic effect of HKE_2_ was likewise inhibited by vatalanib.

Activation of VEGFR2 by HKE_2_ was unlikely to involve direct binding of the lipid as an agonist to the ligand binding domain of the receptor. The latter is designed to recognize a protein ligand that has little to no structural similarity to an oxygenated derivative of arachidonic acid. In fact, binding studies with the soluble extracellular ligand binding domain of human VEGFR2 did not indicate binding of HKE_2_. Thus, HKE_2_ was assumed to activate VEGFR2 indirectly, and two possibilities were explored. We first tested covalent binding of HKE_2_ to matrix metalloproteinase MM9 as a means to activate MMP9 and stimulate release of VEGF from extracellular matrix storage, resulting in activation of VEGFR2 (26, 27). Covalent binding of HKE_2_ to the cysteine thiol ligand that keeps the catalytic Zn^2+^ in an inactive form in MMP9, appeared feasible based on the covalent binding of the electrophilic enone moiety of HKE_2_ with glutathione (4). Adduction of HKE_2_ to MMP9, however, could not be established. We next tested whether HKE_2_ targets and inhibits protein phosphatase PTP1B, the enzyme that de-phosphorylates activated VEGFR2 (29). Using reagents for the detection and analysis of reactive cysteine residues we showed that HKE_2_ covalently binds to PTP1B and that adduction by HKE_2_ inhibited enzymatic activity. Together, our studies suggested that inhibition of PTP1B by HKE_2_ results in decreased de-phosphorylation of VEGFR2, a mechanism consistent with the time course of the effects of HKE_2_ on HUVEC tubulogenesis.

Oxidative modification is a major posttranslational mechanism contributing to the dynamic regulation of protein tyrosine phosphorylation (45). Both protein kinases and phosphatases undergo redox modifications of catalytically crucial cysteine residues by reactive oxygen species like hydrogen peroxide or by small molecule electrophiles, including lipid-derived electrophiles like 4-hydroxy-nonenal or acrolein (30, 46, 47). The latter contain an enone moiety that reacts with a nucleophilic cysteine in a Michael-type reaction resulting in covalent modification of the residue by the electrophile. The phosphatase PTP1B undergoes H_2_O_2_-mediated sulfenylation of the active site cysteine that triggers a conformational change resulting in inhibition of the enzyme (48, 49). Small molecule electrophiles have been described to covalently inhibit various protein phosphatases including PTP1B. Electrophiles were found to bind either to the active site cysteine of PTP1B or to a cysteine at an allosteric site, with both binding events resulting in inhibition of the enzyme (30, 50). Thus, covalent binding of PTP1B by HKE_2_ likely involves a Michael-type reaction of the enone moiety with the catalytic or allosteric cysteine residue of the enzyme and follows an established mechanism of inhibition of this phosphatase.

The initial screen for RTK targets of HKE_2_ in HUVEC also showed enhanced phosphorylation of ALK, EphA10, RYK, and Tie1 (Fig. 1D). Anaplastic lymphoma kinase (ALK) is also a substrate for PTP1B (51), and thus HKE_2_-mediated inhibition of PTP1B may account for the enhanced phosphorylation of ALK. Phosphatases for EphA10 and RYK could not be identified in a literature search. The potential HKE_2_-target Tie1 is dephosphorylated by vascular endothelial protein tyrosine phosphatase (VE-PTP) (52). VE-PTP is a receptor-type PTP that also undergoes redox-regulation at an active site cysteine (53), and therefore, may be inhibited by HKE2 in a mechanism similar to PTP1B, thus possibly explaining the observed HKE2-dependent phosphorylation of Tie1 (Fig. 1D). To take the results from the RTK screen one step further, and considering that a large number of phosphatases and other proteins undergo some form of redox control (54, 55) and thus are potential targets of HKE2, finding only a handful of targeted RTK suggests that HKE2 exerts a particular selectivity regarding which phosphatases (and possibly other proteins) are covalently modified. The unexpectedly small number of putative target proteins when compared to other reactive enone electrophiles (46) may at least in part be due to the high polarity provided by the 4 hydroxyl groups surrounding the enone moiety of HKE2.

The stimulation of angiogenesis and tubulogenesis by HKE_2_ was comparable to the effect achieved by VEGF although it is difficult to directly compare the potencies of a protein and an eicosanoid. Compared to other eicosanoids, HKE_2_ appeared to be more potent. The *in vitro* pro-tubulogenic effect of EETs, for example, required 10-20 times higher concentrations (25). When tested at the same concentration as HKE_2_, PGE_2_ and 5-HETE did not stimulate endothelial cell tubulogenesis. This result was at odds with reported pro-angiogenic properties of 5-HETE and PGE_2_ but may be due to differences in the experimental conditions. It appears that the biosynthetic cross-over pathway using consecutive oxygenation of AA by 5-LOX and COX-2 provides HKE_2_ as a pro-angiogenic eicosanoid that was more potent than the biosynthetic intermediate 5-HETE or PGE_2_ as a prototypical COX-2 derived pro-angiogenic product.

We have shown that HKE_2_ is sufficient to stimulate angiogenesis *in vivo* when exogenously supplied in the sponge model but it is not known whether HKE_2_ is also required during physiological or pathophysiological angiogenesis. In order to determine the activity of endogenously formed HKE_2_ specific tools are necessary to enable manipulation of HK formation without affecting other eicosanoids derived from the 5-LOX and COX-2 biosynthetic pathways. For example, genetic deletion of 5-LOX or COX-2 will eliminate not only HKE_2_ but also a host of other eicosanoids, some of which known to contribute to angiogenesis directly or indirectly. A pharmacologic approach is likewise unsuited since there are no known inhibitors that selectively decrease HK biosynthesis without affecting any of the other eicosanoids derived from either 5-LOX or COX-2. An HK-specific antibody may be an appropriate tool but such a reagent is not available or in development as far as we are aware.

Targeting the VEGF pathway is a mainstay in cancer therapy but treatment is associated with significant adverse effects (56). The potent activation of VEGFR2 signaling by HKE_2_ suggests blocking biosynthesis of the latter as a possible clinical approach to inhibit angiogenesis. If a role of HKE_2_ can be established in pathologic angiogenesis in certain types of cancer or in certain patients then targeting of 5-LOX or COX-2 using approved drugs like 5-LOX inhibitors or NSAIDs, respectively, may offer a way to reduce the dosing of anti-VEGF drugs and thereby decrease the associated side effects. In fact, the usefulness of COX inhibitors in chemoprevention and treatment of some cancers has been well established (57, 58). It remains to be shown whether some of the therapeutic effects of COX inhibition in cancer might be due to preventing formation of the pro-angiogenic HK eicosanoids derived from the cross-over of the 5-LOX and COX-2 enzymes.

## Material and Methods

### Materials

HKE_2_ was prepared by total synthesis as described (10). Other eicosanoids and the VEGFR2 inhibitor vatalanib hydrochloride were obtained from Cayman Chemical. The human phospho-Receptor Tyrosine Kinase array kit was from R&D Systems. Recombinant human protein tyrosine phosphatase 1B (PTP1B) was from Abcam. Trypsin, HBSS, Rhodamine B isothiocyanate-dextran (MW 70 kDa), phorbol 12-myristate 13-acetate (PMA), and bovine serum albumin (BSA) were from Sigma. Fetal bovine serum (FBS), and growth factor reduced matrigel were from Corning (Corning, MA, USA). Medium 200PRF and low serum growth supplement (LSGS) were purchased from Invitrogen (USA). The selective inhibitors U0126 (ERK), SP600125 (JNK), and LY294002 (PI3K), and cell lysis buffer were from Cell Signaling. Vascular endothelial growth factor-165 (VEGF_165_) was purchased from PeproTech (Cranbury, NJ, USA). Ketoprofen was purchased from Zoetis (Kalamazoo, MI, USA). The MMP inhibitor GM6001 was a kind gift from Dr. Barbara Fingleton, Vanderbilt University School of Medicine, Department of Pharmacology. Organic coconut MCT oil containing no trans, mono-, or polyunsaturated fat was purchased from a local supermarket.

### HUVEC and cell culture conditions

Human umbilical vein endothelial cells (HUVEC) and cell culture medium (Vascular Cell Basal Medium completed with Endothelial Cell Growth Kit-VEGF) were obtained from ATCC (Manassas, VA, USA). Cells were cultured at a density between 5000 – 10000 cells/cm^2^ in 10-cm dishes. Culture medium was replaced every other day until 80 – 90% confluence was reached. Passage number was between 3 and 7, and population doubling level (PDL) values between 8 – 11 were used for all the experiments.

### Human phospho-receptor tyrosine kinase array

HUVEC were seeded in 10 cm dishes at a density of 4400 cells/cm^2^ and incubated until 70 - 80% confluent. Cells were then washed twice with HBSS and incubated in Medium 200 PRF. Twelve hours later, the cells were treated with 50, 100, or 500 nM HKE_2_ or vehicle control (ethanol; 0.1% v/v) for 30 min. The dishes were then placed on ice, the cells were washed twice with ice-cold PBS and lysed in lysis Buffer 17 (R&D System) containing 10 μg/mL aprotinin (Sigma), 10 μg/mL leupeptin (Millipore), 10 μg/mL pepstatin (Sigma), and protease inhibitor (Sigma). After incubation at 4°C for 30 min, the cell lysates were centrifuged at 14,000 x g for 5 min, and the supernatant was transferred to a clean 1.5 mL tube. The protein content in the cell lysis supernatant was quantified using the BCA assay. A total amount of 100 μg protein, pooled from 4 independent experiments, was incubated with a phospho-RTK array membrane at 4°C overnight according to the manufacturer’s instructions. On the following day, membranes were washed and probed with anti-phospho-tyrosine-HRP detection antibody at room temperature for 2 h. Membranes were developed with Chemi Reagent Mix prior to exposure to X-ray film and analysis by ImageJ software.

### Western blot

Serum starved HUVEC (∼70-80% confluent) were treated with HKE_2_ (50-500 nM), VEGF_165_ (50 ng/mL), or vehicle (ethanol, 0.1% v/v). After 0.5-2 h cell lysates were analyzed by SDS-PAGE followed by Western blot for levels of phosphorylated tyrosine, phosphorylated and total VEGFR2, phosphorylated and total ERK 1/2, Akt, and p38 MAPK, and β-actin. Primary antibodies used were: phospho-tyrosine (P-Tyr-100), phospho-VEGFR2 (Tyr1175) (19A10), VEGFR2 (55B11), phospho-p44/42 MAPK (Erk1/2) (Thr202/Tyr204) (20G11), p44/42 MAPK (Erk1/2) (137F5), phospho-Akt (Ser473) (D9E) XP^®^, Akt, phospho-p38 MAPK (Thr180/Tyr182) (D3F9) XP^®^ p38 MAPK, β-actin (8H10D10) (all from Cell Signaling Technology), and PTP1B (FG6-1G) (Calbiochem). Membranes were incubated with primary antibody at 4°C overnight, and subsequently incubated with secondary antibody, IRDye 680LT goat anti-rabbit (926-68024) or IRDye 800CW donkey anti-goat (926-32214) (LI-COR Biosciences) (1:10,000 dilution), at room temperature for 1 h. Membranes were scanned by Li-Cor Odyssey Infrared Imaging System. Bands were quantified using ImageJ software. The level of phosphorylated and total protein in each group was expressed as the ratio between phospho-/total-protein. Three or four independent experiments were performed.

### Tubulogenesis assay

The tubulogenesis assay was performed and quantified as previously described (59). For the assay, 96-wells plates were coated with 50 µL reduced growth factor matrigel and incubated at 37°C for 60 min to solidify the gel. HUVEC at 80% confluence were starved for 2 h in Medium 200PRF. Cells were then trypsinized, the concentration adjusted to 1 × 10^5^ cells/mL in Medium 200PRF and 50 µL were added to each well (5 × 10^3^ cells/well). For inhibitor treated cells, 50 µL of 150 nM vatalanib (75 nM final, ethanol; 0.0025% v/v) or vehicle were added and incubated for 2 h. After the pre-incubation, the cells were stimulated with HKE_2_ (10 or 50 nM final, ethanol; 0.02% v/v), VEGF_165_ (50 ng/mL) or vehicle. After 6 hours at 37°C, images were taken at 4x magnification using EVOS XL Core Imaging System (ThermoFisher, USA). One image for each treatment (performed in duplicate) was taken. ImageJ software with the Angiogenesis Analyzer plugin (60, 61) was used to determine tube network formation by measuring contact points between three different cells (junctions), segments delimited between two junctions, number of rings, total number of junctions and segments, and the total area of rings formed. The ImageJ software enables automatic and unbiased scoring of endothelial cell network formation.

### In vivo angiogenesis

Experiments were approved by the Vanderbilt University IACUC (Protocol # M1900044-00), and NIH principles of laboratory animal care were followed. We used the subcutaneous sponge model to determine the effects of HKE_2_ on *in vivo* angiogenesis (25). Two sterile polyvinyl acetal CF-50 round sponges (diameter, 8 mm; thickness, 3 mm; a kind gift from Dr. Beate Eckes, University of Cologne, Germany) were implanted under the dorsal skin of BALB\c female mice (12 weeks of age, 20 g).

Mice were administrated the VEGFR2 inhibitor, vatalanib, via daily gavage (in DMSO/polyethylene glycol 400/0.9% sodium chloride 5/47.5/47.5 (by vol.), 50 mg/kg p.o.). At the same time, the sponges were subcutaneously injected every other day with 50 μL of either MCT (medium-chain triglyceride) oil (vehicle, n = 8; vehicle + inhibitor, n = 4), HKE_2_ (10 µM in MCT oil, n = 12; HKE_2_ + inhibitor, n = 6), or VEGF_165_ (5 µg/mL in PBS, n = 7; VEGF_165_ + inhibitor, n = 5). When both inhibitor and HKE_2_ were injected on the same day, inhibitor was orally administrated 2 h before HKE_2_ injection. After two weeks, the mice were injected intravenously with 100 μL of rhodamine-dextran (Mr 75 kD, 3% in PBS) to label blood vessels (25). After 15 minutes, the sponges were collected and analyzed under an epifluorescence microscope. Rhodamine-dextran-positive structures were imaged, the color images converted to black and white pictures using Photoshop (Adobe), and processed as described (25). Vascularity within sponges was expressed as a percentage of area occupied by rhodamine-dextran-positive structures per microscopic field. Five-six images/sponge per treatment were used for analysis.

### Covalent modification of PTP1B

Recombinant human PTP1B (500 nM) was incubated with 5 or 45 µM of HKE_2_ or NEM for 2 h at 37°C. The samples were then incubated with 0.25 mM of Biotin-maleimide (Thermo Fisher Scientific, USA) for 2 h on ice, followed by SDS-PAGE using a 10% polyacrylamide gel. Protein was transblotted onto a PVDF membrane and incubated with 0.25% polyvinylpyrrolidone (Sigma, St. Louis, MO, USA) in Tris-buffered saline containing 0.1% Tween 20 for blocking, washed, and treated with HRP-linked Avidin for 1 h at room temperature. Derivatized protein bands were detected by addition of Chemi-Lumi One L Western blotting detection reagents (Nacalai Tesque, Tokyo, Japan) and visualized using a WSE-6100 LuminoGraph I (ATTO, Tokyo, Japan). For the band shift assay, samples were incubated with PEG-PC-maleimide (Dojindo, Japan) for 30 min at 37°C, resulting in a shift of the MW of the adducted protein by about 5 kD for every cysteine residue derivatized with the reagent. The samples were electrophoresed using a reduced SDS-PAGE (10% polyacrylamide gel) and the gel was stained with Coomassie Brilliant Blue. Bands were quantified using Image J software.

### PTP1B phosphatase activity

A phosphatase dilution buffer containing 25 mM HEPES pH 7.2, 50 mM NaCl, 2.5 mM EDTA, 5 mM DTT, and 50 ng/µL BSA was used for the hydrolysis of *p*-nitrophenylphosphate (*p*NPP) to quantify activity of recombinant human PTP1B. PTP1B (1 µM) was incubated with 10 or 90 µM of HKE_2_ or NEM for 30 min at 37°C, and then 10 µL of the PTP1B sample was mixed with 50 µL of *p*NPP (1.5 mM) in phosphatase dilution buffer and 40 µL phosphatase dilution buffer in a 96-well plate. A well with an equal volume of phosphatase dilution buffer was used as a blank. The 96-well plate was incubated at 37°C for 20 min, and the reaction was terminated by the addition of 50 μL of 2 M NaOH stopping solution, and the wavelength at 405 nm was measured.

### Statistical analysis

Prism 5.0 (GraphPad, La Jolla, CA, USA) was used for statistical analysis of the data. Student’s t-test or ANOVA followed by Newman-Keuls post-hoc analysis were used for the analysis of normally distributed data. Statistically significant differences were considered at *p < 0*.*05*.

## Acknowledgement

This work was supported by National Institutes of Health grants R01GM076592 and R01GM118412 to C.S., R01CA162433 and R01DK095761 to A. P, and R01GM115722 to G.A.S.. A.P. is also supported by a Veterans Affairs Merit Review 1I01BX002025-01. A.P. is the recipient of a Veterans Affairs Senior Research Career Award. J.A.G.-B. was supported by a postdoctoral award from the American Heart Association (16POST30690001). The project was in part supported by CTSA award No. UL1 TR002243 from the National Center for Advancing Translational Sciences. Use of the Monolith NT MST instrument, located in the Vanderbilt University Computational Structural and Chemical Biology Instrument Core, was supported by NIH grant 1S10OD021483-01. The content is solely the responsibility of the authors and does not necessarily represent the official views of the National Institutes of Health.

## Abbreviations

AKT: protein kinase B
ERK: extracellular signal-regulated kinase
COX: cyclooxygenase
EET: epoxyeicosatrienoic acid
HETE: hydroxyeicosatetraenoic acid
HK: hemiketal
HUVEC: human umbilical vein endothelial cells
LOX: lipoxygenase
MCT: medium-chain triglyceride
MST: microscale thermophoresis
NEM: *N*-ethyl-maleimide
*p*NPP: *p*-nitrophenylphosphate
PTP1B: protein tyrosine phosphatase 1B
RTK: receptor tyrosine kinase
VEGF: vascular endothelial growth factor
VEGFR: vascular endothelial growth factor receptor

## Notes

### Competing Interest Statement

The authors have declared no competing interest.

