## Supplemental Figures for "The 5-LOX/COX-2 cross-over metabolite, hemiketal E_2_, enhances VEGFR2 activation and promotes angiogenesis"

[a]Division of Clinical Pharmacology, Department of Pharmacology, and Vanderbilt Institute of Chemical Biology, [b]Department of Chemistry and Vanderbilt Institute of Chemical Biology, Vanderbilt University Nashville, Tennessee 37232, U.S.A.; [c]Division of Nephrology, Department of Medicine, Vanderbilt University School of Medicine, Nashville, Tennessee 37232, U.S.A.; [d]Graduate School of Bioagricultural Sciences, Nagoya University, Nagoya, 464-8601, Japan; [e]Veterans Affairs Hospital, Nashville, TN, 37232, U.S.A.

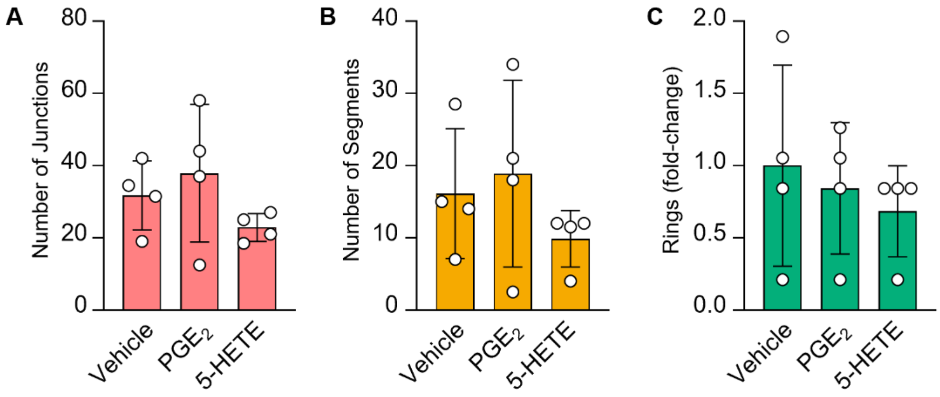


**Fig. S1.** Effect of $PGE_2$ and 5*S*-HETE on HUVEC tubulogenesis. Quantification of (A) junctions, (B) segments, and (C) rings in HUVEC treated with ethanol (vehicle), $PGE_2$ or 5*S*-HETE (100 nM each). After 6 hours, capillary network formation was visualized and quantified by ImageJ software with the Angiogenesis Analyzer plugin counting the number of (A) junctions, (B) segments, and (C) rings. Data are representative of 3 independent experiments performed in duplicate. Data are presented as means ±S.D. No significant difference was observed.

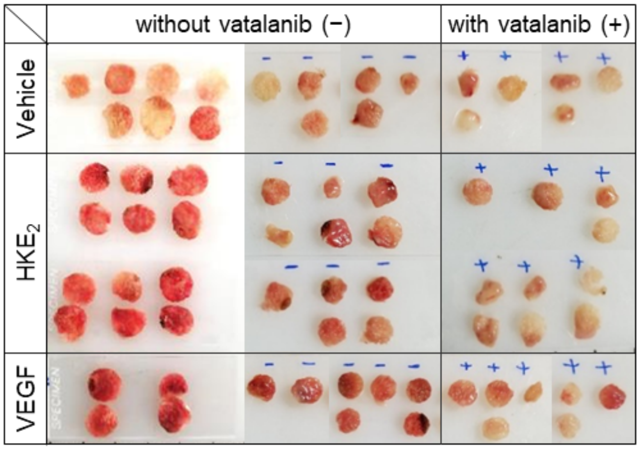


**Fig. S2.** *In vivo* angiogenesis induced by $HKE_2$. Gross images of polyvinyl acetal CF-50 sponges implanted under the dorsal skin of BALB\c female mice (12 weeks of age, 20 g) treated with vehicle (MCT oil; 50 μL), $HKE_2$ (10 μM; 50 μL), or $VEGF_{165}$ (5 μg/mL; 50 μL) with (+) or without (-) oral administration of the VEGFR2 inhibitor vatalanib for 2 weeks. Sponges derived from the same mouse are placed vertically. Some of the implanted sponges were lost during treatment and were not included in the analysis. Three independent experiments were performed. The total number of mice used for each group was: Vehicle; n = 8, vehicle + vatalanib; n = 4, $HKE_2$; n = 12, $HKE_2$ + vatalanib; n = 6, $VEGF_{165}$; n = 7, and $VEGF_{165}$ + vatalanib; n = 5.

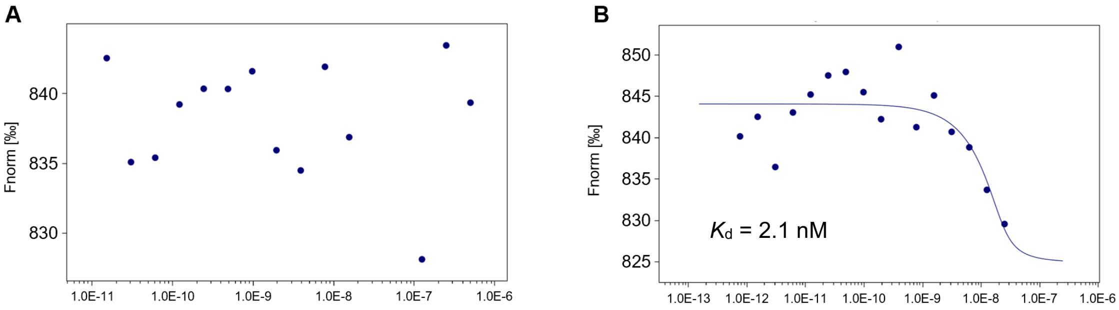


**Fig. S3.** HKE$_2$ does not bind to the ligand binding domain of human VEGFR2. The biotinylated extracellular domain of human VEGFR2 (NP_002244.1) (Met$_1$-Glu$_{764}$) fused with a polyhistidine tag at the C-terminus (10012-H08H-B, Sinobiological) was labeled using Monolith Protein Labeling Kit RED-NHS 2$^{nd}$ Generation (MONOTEMPER) according to the manufacturer's instructions. As a ligand, 25 nM VEGF$_{165}$ or 500 nM HKE$_2$ were prepared in PBS containing 0.005% Tween 20, and 10-fold serial dilutions were made 15 times. The same volumes of two-times concentrated RED-labeled VEGFR2 (430 nM VEGFR2) in PBS containing 0.005% Tween 20 and 1.5 pM-25 nM VEGFR$_{165}$ or 30 pM-500 nM HKE$_2$ were mixed to prepare 215 nM RED-labeled VEGFR2 for microscale thermophoresis (MST). Samples were infused into Monolith NT 115 capillaries and MST measurements were performed using a Monolith NT.115 instrument (201602-BR-NO13). The data points for HKE$_2$ concentration of 62.5 and 31.3 nM are missing due to incomplete filling of the capillaries. For HKE$_2$ a dissociation constant ($K_d$) could not be calculated.

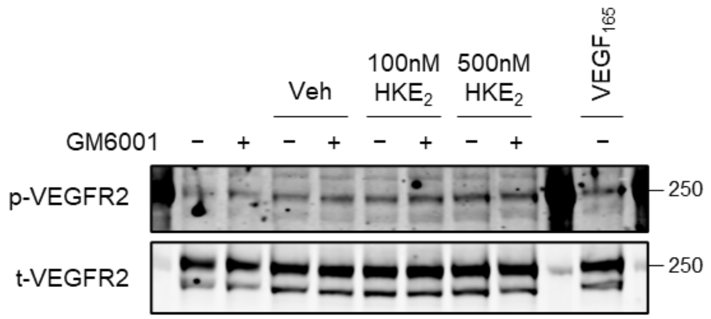


**Fig. S4.** $HKE_2$ does not activate MMP9. Serum starved HUVEC (~70-80% confluent) were pre-treated with 25 μM of the MMP inhibitor GM6001 for 1 h followed by $HKE_2$ (100 or 500 nM), $VEGF_{165}$ (50 ng/mL), or vehicle (ethanol, 0.1% v/v). After 30 min cell lysates were analyzed by SDS-PAGE followed by Western blot for levels of phospho-VEGFR2 (Tyr1175) (19A10), and VEGFR2 (55B11). Membranes were incubated with primary antibody at 4°C overnight, and subsequently incubated with secondary antibody, IRDye 680LT goat anti-rabbit (926-68024) (LI-COR Biosciences) (1:10,000 dilution), at room temperature for 1 h. Membranes were scanned by Li-Cor Odyssey Infrared Imaging System.
